# Efficient Proximal Gradient Algorithm for Inference of Differential Gene Networks

**DOI:** 10.1101/450130

**Authors:** Chen Wang, Feng Gao, Georgios B. Giannakis, Gennaro D’Urso, Xiaodong Cai

**Affiliations:** Department of Electrical and Computer Engineering, University of Miami, Coral Gables, Florida 33146, United States; Department of Electrical and Computer Engineering, University of Minnesota, 55455 Minneapolis, MN; Department of Molecular and Cellular Pharmacology, University of Miami, 33136 Miami, FL; Sylvester Comprehensive Cancer Center, University of Miami, Miami, Florida 33136, United States

**Keywords:** Gene network, differential network, proximal gradient method

## Abstract

**Background:** Gene networks in living cells can change depending on various conditions such as caused by different environments, tissue types, disease states, and development stages. Identifying the differential changes in gene networks is very important to understand molecular basis of various biological process. While existing algorithms can be used to infer two gene networks separately from gene expression data under two different conditions, and then to identify network changes, such an approach does not exploit the data jointly, and it is thus suboptimal. A desirable approach would be clearly to infer two gene networks jointly, which can yield improved estimates of network changes.

**Results:** In this paper, we developed a proximal gradient algorithm for differential network (ProGAdNet) inference, that jointly infers two gene networks under different conditions and then identifies changes in the network structure. Computer simulations demonstrated that our ProGAdNet outperformed existing algorithms in terms of inference accuracy, and was much faster than a similar approach for joint inference of gene networks. Gene expression data of breast tumors and normal tissues in the TCGA database were analyzed with our ProGAdNet, and revealed that 268 genes were involved in the changed network edges. Gene set enrichment analysis of this set of 268 genes identified a number of gene sets related to breast cancer or other types of cancer, which corroborated the gene set identified by ProGAdNet was very informative about the cancer disease status. A software package implementing the ProGAdNet and computer simulations is available upon request.

**Conclusion:** With its superior performance over existing algorithms, ProGAdNet provides a valuable tool for finding changes in gene networks, which may aid the discovery of gene-gene interactions changed under different conditions.

## Background

Genes in living cells interact and form a complex network to regulate molecular functions and biological processes. Gene networks can undergo toplogical changes depending on the molecular context in which they operate [1, 2]. For example, it was observed that transcription factors (TFs) can bind to and thus regulate different target genes under varying environmental conditions [3, 4]. Changes of genetic interactions when cells are challenged by DNA damage as observed in [5] may also reflect the structural changes of the underlying gene network. This kind of rewiring of gene networks has been observed not only in yeast [3–6], but also in mammalian cells [7, 8]. More generally, differential changes of gene networks can occur depending on environment, tissue type, disease state, development and speciation [1]. Therefore, identification of such differential changes in gene networks is of paramount importance when it comes to understanding the molecular basis of various biological processes.

Although a number of computational methods have been developed to infer the structure of gene regulatory networks from gene expression and related data [9-12], they are mainly concerned with the static structure of gene networks under a single condition. These methods rely on similarity measures such as the correlation or mutual information [13, 14], Gaussian graphical models (GGMs) [15, 16], Bayesian networks [17, 18], or linear regression models [19-22]. Existing methods for the analysis of *differential* gene interactions under different conditions typically attempt to identify differential co-expression of genes based on correlations between their expression levels [23]. While it is possible to use an existing method to infer a gene network under different conditions separately, and then compare the inferred networks to determine their changes, such an approach does not jointly leverage the data under different conditions in the inference; thus, it may markedly sacrifice the accuracy in the inference of network changes.

In this paper, we develop a very efficient proximal gradient algorithm for differential network (ProGAdNet) inference, that jointly infers gene networks under two different conditions and then identifies changes in these two networks. To overcome the challenge of the small sample size and a large number of unknowns, which is common to inference of gene networks, our method exploits two important attributes of gene networks: i) sparsity in the underlying connectivity, meaning that the number of gene-gene interactions in a gene network is much smaller than the number of all possible interactions [19, 24-26]; and, ii) sparsity in the structural changes, meaning that the number of interactions changed in response to different conditions is much smaller than the total number of interactions present in the network. A similar network inference setup was considered in [27] for inferring multiple gene networks, but no new algorithm was developed there; instead [27] adopted the lqa algorithm of [28] that was designed for generalized linear models. Our computer simulations demonstrated superior performance of our ProGAdNet algorithm relative to existing methods including the lqa algorithm. Analysis of a set of RNA-Seq data from normal tissues and breast tumors with ProGAdNet identified genes involved in changes of the gene network.

The differential gene-gene interactions identified by our ProGAdNet algorithm yield a list of genes alternative to the list of differentially expressed genes. This may provide additional insight into the molecular mechanism behind the phenotypical difference of the tissue under different conditions. Alternatively, the two gene networks inferred by our ProGAdNet algorithm can be used for further differential network analysis (DiNA). DiNA has received much attention recently; the performance often DiNA algorithms was assessed in [29] using gene networks and gene/microRNA networks. Given two networks with the same set of nodes, a DiNA algorithm computes a score for each node based on the difference of global and/or local topologies of the two networks, and then ranks nodes based on these scores. Apparently, DiNA relies on the two networks that typically need to be constructed from certain data. Our ProGAdNet algorithm provides an efficient and effective tool for constructing two gene networks of the same set of genes from gene expression data under two different conditions, which can be used by a DiNA algorithm for further analysis.

## Methods

### Gene Network Model

Suppose that expression levels of *p* genes have been measured with microarray or RNA-seq, and let *X_i_* be the expression level of the ith gene, where *i* = 1, …, *p*. To identify the possible regulatory effect of other genes on the ith gene, we employ the following linear regression model as also used in [19-22]

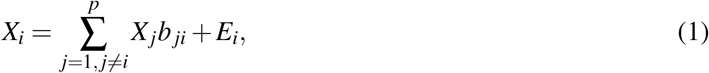

where *E_i_* is the error term, and unknown regression coefficients (*b_ji_*)’s reflect the correlation between *X_i_* and *X_j_* after adjusting the effects of other variables, 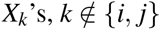. This adjusted correlation may be the result of possible interaction between genes *i* and *j*. The nonzero (*b_ji_*)’s define the edges in the gene network. Suppose that *n* samples of gene expression levels of the same organism (or the same type of tissue of an organism) under two different conditions are available, and let *n* × 1 vectors **x**_i_ and 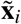 contain these *n* samples of the *i*th gene under two conditions, respectively. Define *n* × *p* matrices 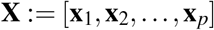 and 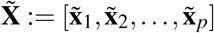, and *p* × *p* matrices **B** and 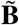 whose element on the *i*th column and the *j*th row are *b_ji_* and 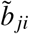, respectively. Letting 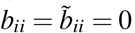, model (1) yields the following

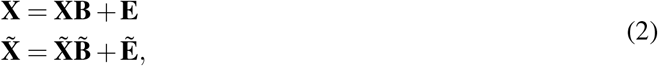

where *n* × *p* matrices **E** and 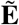 contain error terms. Matrices **B** and 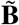 characterize the structure of the gene networks under two conditions.

Our main goal is to identify the changes in the gene network under two conditions, namely, those edges from gene *j* to gene *i* such that 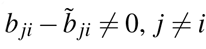. One straightforward way to do this is to estimate **B** and 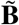 separately from two linear models in (2), and then find gene pairs (*i, j*) for which 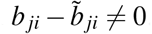. However,this approach may not be optimal, since it does not exploit the fact that the network structure does not change significantly under two conditions, that is, most entries of **B** and 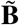 are identical. A better approach is apparently to infer gene networks under two conditions jointly, which can exploit the similarity between two network structures and thereby improve the inference accuracy.

If we denote the *i*th column of **B** and 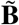 as **b**_i_ and 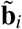, we can also write model (2) for each gene separately as follows: **x_*i*_** = **Xb_*i*_** + **e_*i*_** and 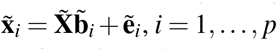, where **e**_*i*_ and 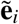 are the *i*th column of **E** and 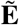, respectively. To remove the constraints *b_ii_* = 0, *i* = 1, …, *p*, we define matrices 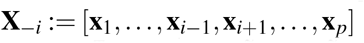 and 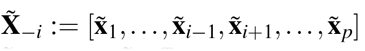, vectors 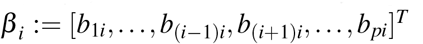 and 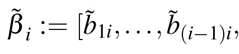 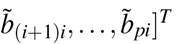. The regression model for the gene network under two conditions can be written as

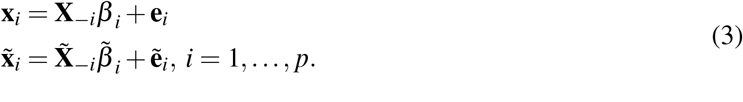

Based on (3), we will develop a proximal gradient algorithm to infer *β_i_* and 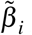 jointly, and identify changes in the network structure.

## Network inference

### Optimization Formulation

As argued in [30-32], gene regulatory networks or more general biochemical networks are sparse, meaning that a gene directly regulates or is regulated by a small number of genes relative to the total number of genes in the network. Taking into account sparsity, only a relatively small number of entries of **B** and 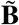, or equivalently entries of *β_i_* and 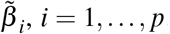, are nonzero. These nonzero entries determine the network structure and the regulatory effect of one gene on other genes. As mentioned earlier, the gene network of an organism is expected to have similar structure under two different conditions. For example, the gene network of a tissue in a disease (such as cancer) state may have changed, comparing to that of the same tissue under the normal condition, but such change in the network structure is expected to be small relative to the overall network structure. Therefore, it is reasonable to expect that the number of edges that change under two conditions is small comparing with the total number of edges of the network.

Taking into account sparsity in the network and also in the changes of the network under two conditions, we formulate the following optimization problem to jointly infer gene networks under two conditions:

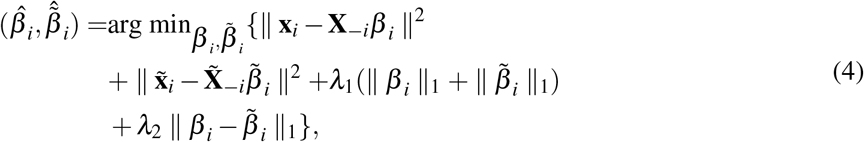

where || · || stands for Euclidean norm, || · ||_1_ stands for *l_1_* norm, and *λ*_1_ and *λ*_2_ are two positive constants. The objective function in (4) consists of the squared error of the linear regression model (1) and two regularization terms 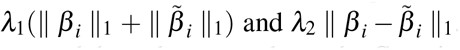. Note that unlike the GGM, the regularized least squared error approach here does not rely on the Gaussian assumption. The two regularization terms induce sparsity in the inferred networks and network changes, respectively. This optimization problem is apparently convex, and therefore it has a unique and globally optimal solution. Note that the term 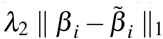 is reminiscent of the fused Lasso [33]. However, all regression coefficients in the fused Lasso are essentially coupled, whereas here the term 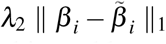 only couples each pair of regression coefficients, *β_ij_* and 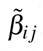 As will be described next, this enables us to develop an algorithm to solve optimization problem (4) that is different from and more efficient than the algorithm for solving the general fused Lasso problem. Note that an optimization problem similar to (4) was formulated in [27] for inferring multiple gene networks, but no new algorithm was developed, instead the problem was solved with the lqa algorithm [28] that was developed for general penalized maximum likelihood inference of generalized linear models including the fused Lasso. Our computer simulations showed that our algorithm not only is much faster than the lqa algorithm, but also yields much more accurate results.

### Proximal Gradient Solver

Define 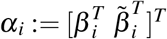 and let us separate the objective function in (4) into the differentiable part *g_1_* (*α_i_*) and the non-differentiable part *g_2_*(*α_i_*) given by

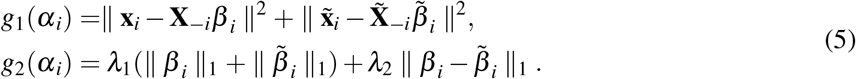

Applying the proximal gradient method [34] to solve the optimization problem (4), we obtain an expression for *α_i_* in the *r*th step of the iterative procedure as follows:

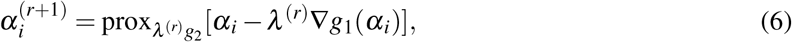

where prox stands for the proximal operator defined as 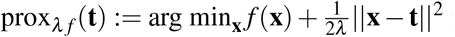 for function *f* (**x**) and a constant vector **t**, and ∇*g_1_* (*α_i_*) is the gradient of *g_1_* (*α_i_*). Generally, the value of step size *λ^(r)^* can be found using a line search step, which can be determined from the Lipschitz constant [34]. For our problem, we will provide a closed-form expression for *λ^(r)^* later. Since *g_1_* (*α_i_*) is simply in a quadratic form, its gradient can be obtained readily as 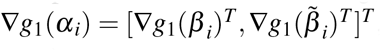, where 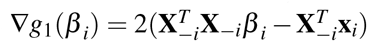 and 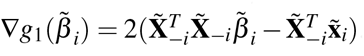.

Upon defining **t** = *β_i_* − *λ*^(*r*)^ ∇*g_1_*(*β_i_*) and 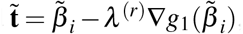, the proximal operator in (6) can be written as

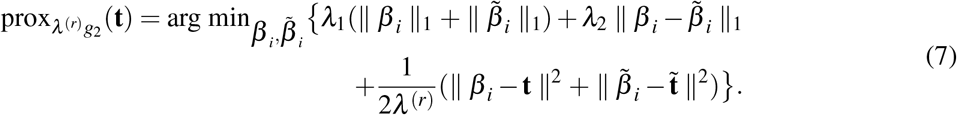

It is seen that the optimization problem in proximal operator (7) can be decomposed into *p* − 1 separate problems as follows

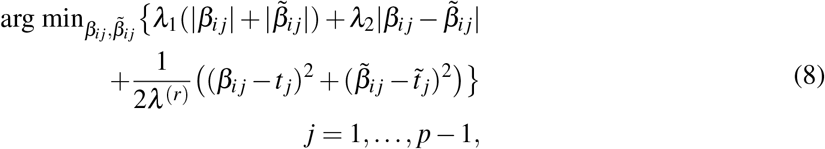

where *β_ij_* and 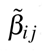 are the *j*th element of *β_i_* and 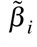, respectively, and *t_j_* and 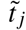 are the *j*th element of **t** and 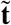, respectively. The optimization problem (8) is in the form of the fused Lasso signal approximator (FLSA) [35]. The general FLSA problem has many variables, and numerical optimization algorithms were developed to solve the FLSA problem [35, 36]. However, our problem has only two variables, which enables us to find the solution of (8) in closed form. This is then used in each step of our proximal gradient algorithm for network inference.

Let us define a soft-thresholding operator *S*(*x,a*) as follows

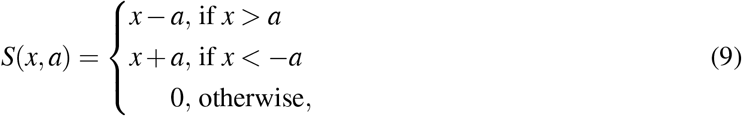

where *a* is a positive constant. Then as shown in [35], if the solution of (8) at *λ*_1_ = 0 is 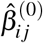 and 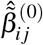, the solution of (8) at *λ*_1_ > 0 is given by

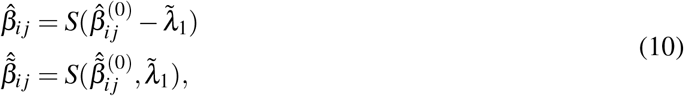

where 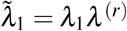. Therefore, if we can solve the problem (8) at *λ*_1_ = 0, we can find the solution of (8) at any *λ*_1_ > 0 from (10). It turns out that the solution of (8) at *λ*_1_ = 0 can be found as

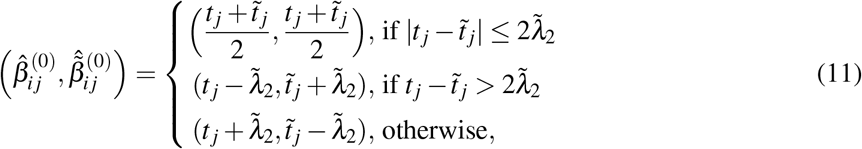

where 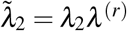. Therefore, our proximal gradient method can solve the network inference problem (6) efficiently through an iterative process, where each step of the iteration solves the optimization problem (6) in closed form specified by (10) and (11). To obtain a complete proximal gradient algorithm, we need to find the step size *λ^(r)^* as will be described next.

### Stepsize

As mentioned in [34], if the step size 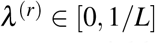, where *L* is the Lipschitz constant of ∇*g_1_* (*α_i_*), then the proximal gradient algorithm converges to yield the optimal solution. We next derive an expression for *L.* Specifically, we need to find *L* such that 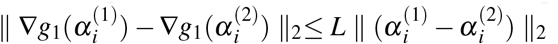, for any 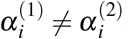, which is equivalent to

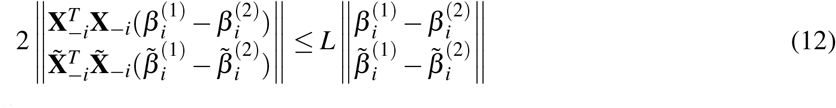

for any 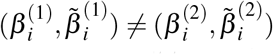 Let *γ* and 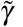 be the maximum eigenvalues of 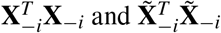, respectively. It is not difficult to see that (12) will be satisfied if 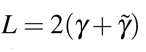. Note that 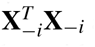 and 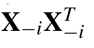 have the same set of eigenvalues. And thus, *γ* can be found using a numerical algorithm with a computational complexity of *O*((min(*n, p*))^2^). After obtaining *L*, the step size of our proximal gradient algorithm can be chosen to be *λ*^(*r*)^ = 1/*L*. Note that *λ*^(*r*)^ does not change across iterations, and it only needs to be computed once. Since the sum of the eigenvalues of a matrix is equal to the trace of matrix, another possible value for *L* is 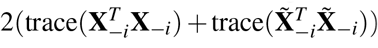, which can save the cost of computing *γ* and 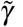. However, this value of *L* is apparently greater than 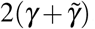, which reduces the step size *λ*^(*r*)^, and may affect the convergence speed of the algorithm.

### Algorithm

The proximal gradient solver of (4) for inference of differential gene networks is abbreviated as ProGAdNet, and is summarized in the following table.

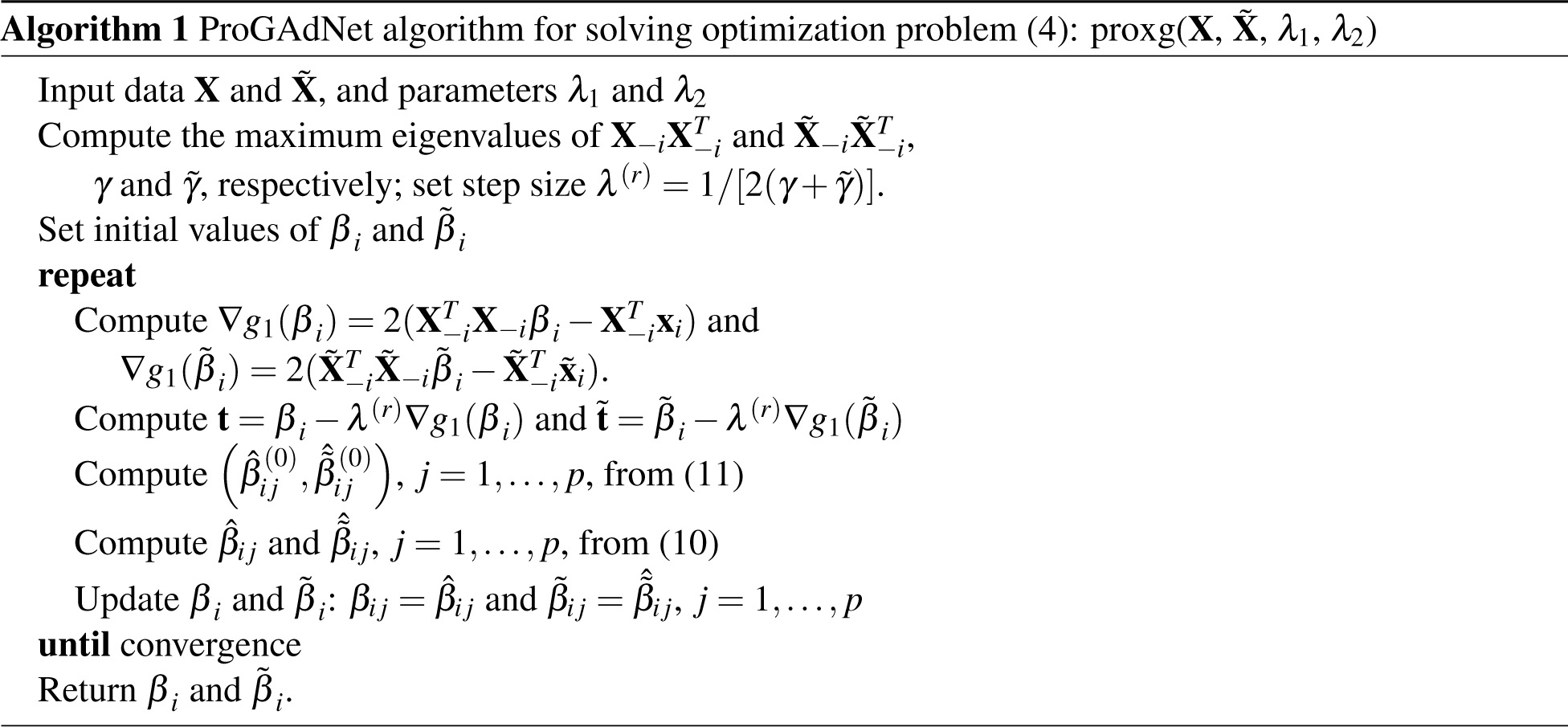

### Maximum values of *λ*_1_ and *λ_2_*

The ProGAdNet solver of (4) is outlined in Algorithm 1 with a specific pair of values of *λ*_1_ and *λ*_2_. However, we typically need to solve the optimization problem (4) over a set of values of *λ*_1_ and *λ*_2_, and then either use cross validation to determine the optimal values of *λ*_1_ and *λ*_2_, or use the stability selection technique to determine nonzero elements of *β_i_* and 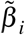, as will be described later. Therefore, we also need to know the maximum values of *λ*_1_ and *λ*_2_. In the following, we will derive expressions for the maximum values of *λ*_1_ and *λ*_2_.

When we determine the maximum values of *λ*_1_, *λ*_1 max_, *λ*_2_ can be omitted in our optimization problem, since when *λ*_1_ = *λ*_1 max_, we have *β_ij_* = 0 and 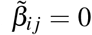, ∀*i* and *j*. Thus, we can use the same method for determining the maximum value of *λ* in the Lasso problem [37] to find *λ*_1 max_, which leads to

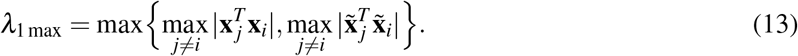

The maximum value of *λ*_2_, *λ*_2 max_ depends on *λ*_1_. It is difficult to find *λ*_2 max_ exactly. Instead, we will find an upper-bound for *λ*_2 max_. Let us denote the objective function in (4) as 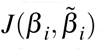, and let the jth column of 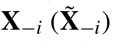 be 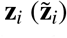. If the optimal solution of (4) is 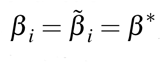, then the subgradient of 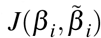 at the optimal solution should contain the zero vector, which yields

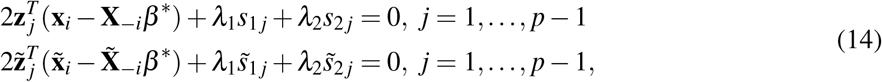

where *s*_1*j*_ = 1 if *β_ij_* > 0, = − 1 if *β_ij_* < 0, or ∈ [−1,1] if *β_ij_* = 0, and *s*_2*j*_ ∈ [−1,1], and similarly, 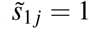 if 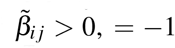 if 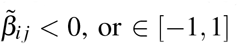 if 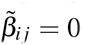, and 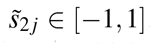. Therefore, we should have *λ*_2_ > 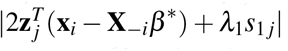 and 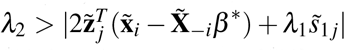, which can be satisfied if we choose *λ*_2_ = 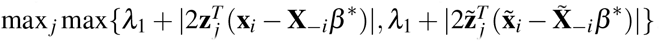. Therefore, the maximum value of *λ*_2_ can be written as

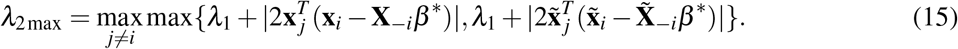

To find *λ*_2 max_ from (15), we need to know *β*\*. This can be done by solving the Lasso problem that minimizes 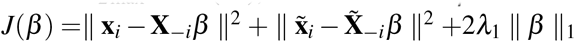 using an efficient algorithm such as glmnet [38].

### Stability Selection

As mentioned earlier, parameter *λ*_1_ encourages sparsity in the inferred gene network, while *λ*_2_ induces sparsity in the changes of the network under two conditions. Generally, larger values of *λ*_1_ and *λ_2_* induce a higher level of sparsity. Therefore, appropriate values of *λ*_1_ and *λ_2_* need to be determined, which can be done with cross validation [38]. However, the nonzero entries of matrices **B** and 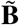, estimated with a specific pair of values of *λ*_1_ and *λ*_2_ determined by cross validation, may not be stable in the sense that small perturbation in the data may result in considerably different **B** and 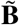. We can employ an alternative technique, named stability selection [39], to select stable variables, as described in the following.

We first determine the maximum value of *λ*_1_, namely *λ*_1 max_, using the method described earlier, then choose a set of *k*,_1_ values for *λ*_1_, denoted as 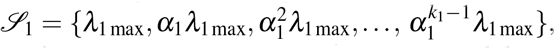, where 0 < *α*_1_ < 1. For each value 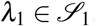, we find the maximum value of *λ*_2_, namely *λ*_2 max_(*λ*_1_), and then choose a set of *k*_2_ values for *λ*_2_, denoted 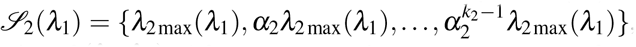, where 0 < *α*_2_ < 1. This gives a set of *K* = *k*_1_ *k*_2_ pairs of (*λ*_1_, *λ*_2_). After we create the parameter space, for each (*λ*_1_, *λ*_2_) pair in the space, we randomly divide the data 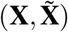 into two subsets of equal size, and infer the network with our proximal gradient algorithm using each subset of the data. We repeat this process for *N* times, which yields 2*N* estimated network matrices, 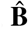 and 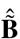. Typically, *N* = 50 is chosen.

Let 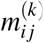, 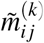, and 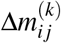 be the number of nonzero 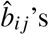 and 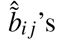, and 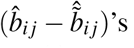, respectively, obtained with the *k*th pair of (*λ*_1_,*λ*_2_). Then, 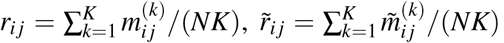 and Δ*r*_ij_ = 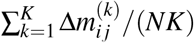 give the frequency of an edge from gene *j* to gene *i* detected under two conditions, and the frequency of the changes for an edge from gene *j* to gene *i,* respectively. A larger *r_ij_*, 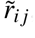 or Δ*r_ij_* indicates a higher likelihood that an edge from gene *j* to gene *i* exists, or the edge from gene *j* to gene *i* has changed. Therefore, we will use *r_ij_* 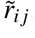 and Δ*r_ij_* to rank the reliability of the detected edges and the changes of edges, respectively. Alternatively, we can declare an edge from gene *j* to gene *i* exists if *r_ij_* ≥ *c* or 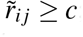; and similarly the edge between gene *j* to gene *i* has changed if Δ*r_ij_* ≥ *c*, where *c* is constant and can be any value in [0.6,0.9] [39].

### Software glmnet and lqa

Two software packages, glmnet and lqa, were used in computer simulations. The software glmnet [38] for solving the Lasso problem is available at https://cran.r-project.org/web/packages/glmnet. The software lqa [28] used in [27] for inferring multiple gene networks is available at https://cran.r-project.org/web/packages/lqa/.

## Results and Discussion

### Computer Simulation with Linear Regression Model

We generated data from one of *p* pairs of linear regression models in (3) instead of all *p* pairs of simultaneous equations in (2), or equivalently (3), as follows. Without loss of generality, let us consider the first equation in (3). The goal was to estimate *β*_1_ and 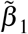, and then identify pairs 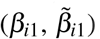, where 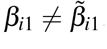. Entries of *n* × (*p* − 1) matrices **X**_−1_ and 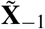 were generated independently from the standardized Gaussian distribution. In the first simulation setup, we chose *n* = 100 and *p* − 1 = 200. Twenty randomly selected entries of *β*_1_ were generated from a random variable uniformly distributed over the intervals [0.5,1.5] and [-1.5,-0.5], and remaining entries were set to zero; 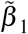 was generated by randomly changing 10 entries of *β*_1_ as follows: 4 randomly selected nonzero entries were set to zero, and 6 randomly selected zero entries were changed to a value uniformly distributed over the intervals [0.5,1.5] and [−1.5, −0.5]. The noise vectors **e**_1_ and 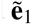 were generated from a Gaussian distribution with mean zero and variance σ^2^ varying from 0.01 to 0.05, 0.1, and 0.5, and then **x**_1_ and **x**_1_ were calculated from (3).

Simulated data **x**_1_, 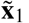, **X**_−1_ and 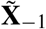 were analyzed with our ProGAdNet, lqa [28] and glmnet [38]. Since lqa was employed by [27], the results of lqa represent the performance of the network inference approach in [27]. The glmnet algorithm implements the Lasso approach in [40]. Both ProGAdNet and lqa estimate *β*_1_ and 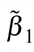 jointly by solving the optimization problem (4), but glmnet estimates *β*_1_ and 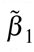 separately, by solving the following two problems separately, 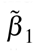 and 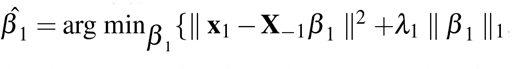 and 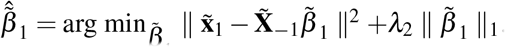. The lqa algorithm uses a local quadratic approximation of the nonsmooth penalty term [41] in the objective function, and therefore, it cannot shrink variables to zero exactly. To alleviate this problem, we set 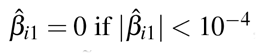, and similarly 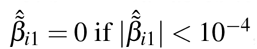, where 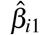 and 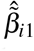 represent the estimates of *β*_*i*1_ and 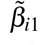, respectively. Five fold cross validation was used to determine the optimal values of parameters *λ*_1_ and *λ*_2_ in the optimization problem. Specifically, for ProGAdNet and lqa, the prediction error (PE) was estimated at each pair of values of *λ*_1_ and *λ*_2_, and the smallest PE along with the corresponding values of *λ*_1_ and *λ*_2_, *λ*_1 min_ and *λ*_2 min_, were determined. Then, the optimal values of *λ*_1_ and *λ*_2_ were the values corresponding to the PE that was two standard error (SE) greater than the minimum PE, and were greater than *λ*_1_ min and A2min, respectively. For glmnet, the optimal values of *λ*_1_ and *λ*_2_ were determined separately also with the two-SE rule.

The inference process was repeated for 50 replicates of the data, and the detection power and the false discovery rate (FDR) for 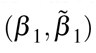 and 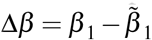 calculated from the results of the 50 replicates in the first simulation setup are plotted in Figure 1. It is seen that all three algorithms offer almost identical power equal or close to 1, but exhibit different FDRs. The FDR of lqa is the highest, whereas the FDR of ProGAdNet is almost the same as that of glmnet for *β*_1_ and 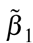, and the lowest for Δ*β*_1_.

**Figure 1.**
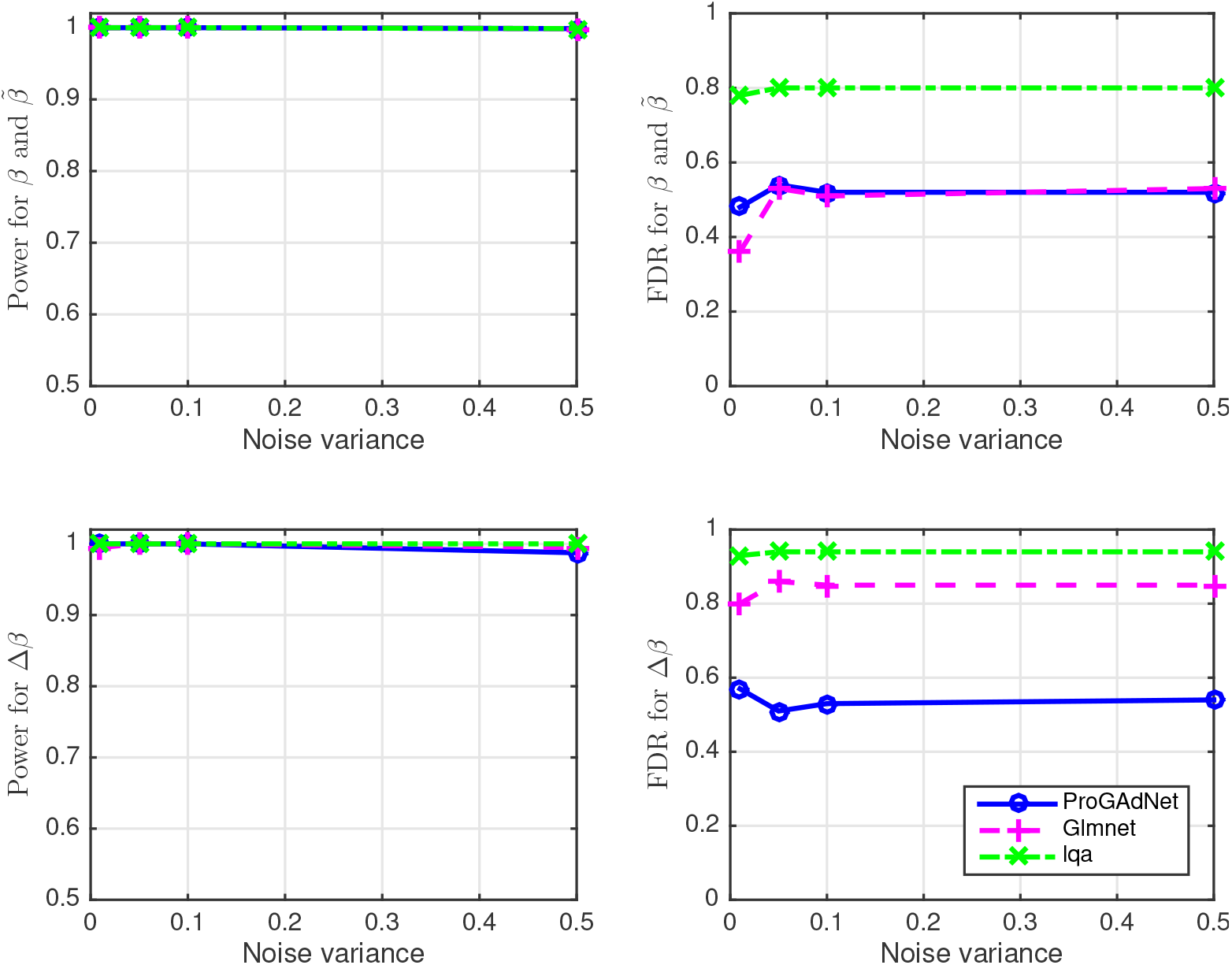
Performance of ProGAdNet, lqa, and Lasso in the inference of linear regression models. Number of samples *n* = 100, and number of variables *p* – 1 = 200.

In the second simulation setup, we let sample size *n* = 150, noise variance σ^2^ = 0.1, and the number of variables *p* – 1 be 500, 800, and 1,000. Detection power and FDR are depicted in Figure 2. Again, the three algorithms have almost identical power, and ProGAdNet offers an FDR similar to that of glmnet, but lower than that of lqa for *β*_1_ and 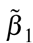, and the lowest FDR for Δ*β*_1_. Simulation results in Figures 1 and 2 demonstrate that our ProGAdNet offers the best performance when compared with glmnet and lqa. The CPU times of one run of ProGAdNet, lqa, and glmnet for inferring a linear model with *n* = 150, *p* – 1 = 1,000, and σ^2^ = 0.1 at the optimal values of *λ*_1_ and *λ*_2_ were 5.82, 145.15, and 0.0037 seconds, respectively.

**Figure 2.**
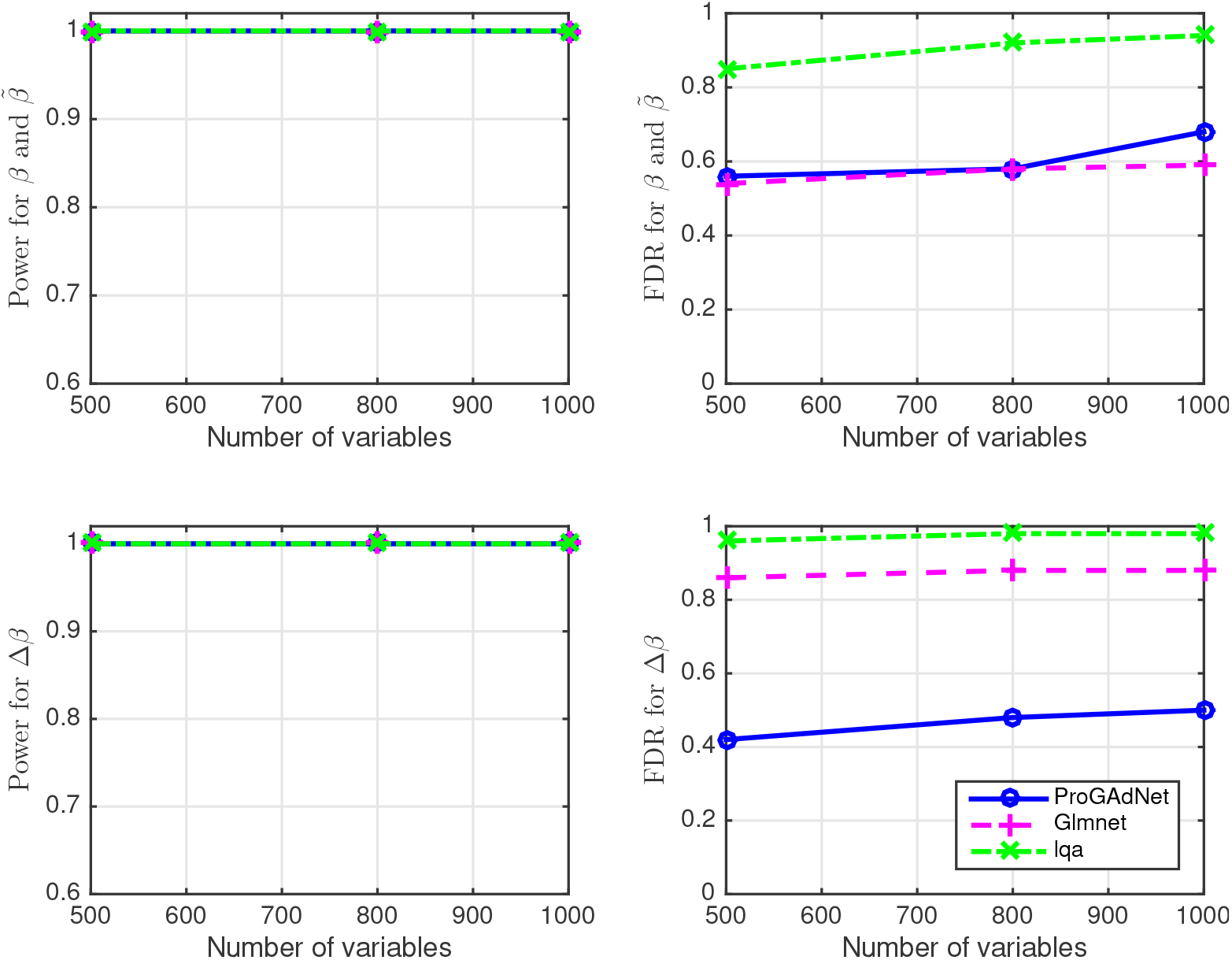
Performance of ProGAdNet, lqa, and Lasso in the inference of linear regression models. Number of samples *n* = 150 and noise variance σ^2^ = 0.1.

Note that although ProGAdNet and lqa solve the same optimization problem, ProGAdNet significantly outperforms lqa. The performance gap is due to the fact that the lqa algorithm uses an approximation of the objective function, whereas our algorithm solves optimization problem (4) exactly. In other words, our ProGAdNet algorithm can always find the optimal solution to the optimization problem, since the objective function is convex, but the lqa algorithm generally cannot find the optimal solution. Moreover, our computer simulations show that our ProGAdNet algorithm is much faster than the lqa algorithm.

### Computer Simulation with Gene Networks

The GeneNetWeaver software [42] was used to generate gene networks whose structures are similar to those of real gene networks. Note that GeneNetWeaver was also employed by the DREAM5 challenge for gene network inference to simulate golden standard networks [12]. GeneNetWeaver outputs an adjacency matrix to characterize a specific network structure. We chose the number of genes in the network to be *p* = 50, and obtained a *p* × *p* adjacency matrix **A** through GeneNetWeaver. The number of nonzero entries of **A**, which determined the edges of the network, was 62. Hence the network is sparse, as the total number of possible edges is *p*(*p* – 1) = 2,450. We randomly changed 6 entries of **A** to yield another matrix **Ã** as the adjacency matrix of the gene network under another condition. Note that the number of changed edges is small relative to the number of existing edges.

After the two network topologies were generated, the next step was to generate gene expression data. Letting *a_ij_* be the entry of **A** on the *i*th row and the *j*th column, we generated a *p* × *p* matrix **B** such that *b_ij_* = 0 if *a_ij_* = 0, and *b_ij_* was randomly sampled from a uniform random variable on the intervals [–1, 0) and (0,1] if *a_ij_* ≠ 0. Another *p* x *p* matrix 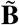 was generated such that 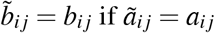, or 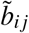 was randomly generated from a uniform random variable on the intervals [–1, 0) and (0,1] if *ã_ij_* ≠ *a_ij_*. Note that (2) gives **X** = (**I – B**)^−1^ **E** and 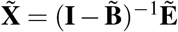. These relationships suggest generating first entries of **E** and 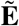 independently from a Gaussian distribution with zero mean and unit variance, and then finding matrices **X** and 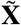 from these two equations, respectively. With real data, gene expression levels **X** and 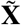 are measured with techniques such as microarray or RNA-seq, and there are always measurement errors. Therefore, we simulated measured gene expression data as **Y** = **X** + **V** and 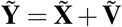, where **V** and 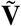 model measurement errors that were independently generated from a Gaussian distribution with zero mean and variance *σ*^2^ that will be specified later. Fifty pairs of network replicates and their gene expression data were generated independently.

Finally, gene networks were inferred with our ProGAdNet algorithm by solving the optimization problem (4), where 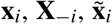, and 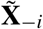 were replaced with the measured gene expression data 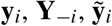, and 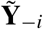. Stability selection was employed to rank the edges that were changed under two conditions. As comparison, we also used Lasso to infer the network topology under each condition by solving the following optimization problems

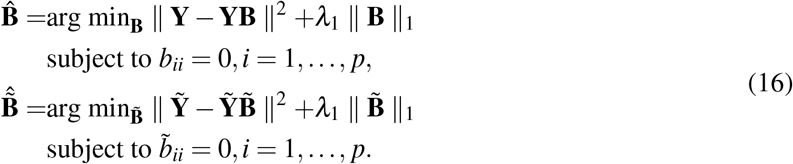

Note that each optimization problem can be decomposed into *p* separate problems that can be solved with Lasso. The glmnet algorithm [38] was again used to implement Lasso. The stability selection technique was employed again to rank the differential edges detected by Lasso. The lqa algorithm was not considered to infer simulated gene networks, because it is very slow and its performance is worse than ProGAdNet and Lasso as shown in the previous section. We also employed the GENIE3 algorithm in [43] to infer **B** and 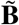 separately, because GENIE3 gave the best overall performance in the DREAM5 challenge [12]. Finally, following the performance assessment procedure in [12], we used the precision-recall (PR) curve and the area under the PR curve (AUPR) to compare the performance of ProGAdNet with that of Lasso and GENIE3. For ProGAdNet and Lasso, the estimate of 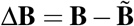 was obtained, and the nonzero entries of Δ**B** were ranked based on their frequencies obtained in stability selection. Then, the PR curve for changed edges was obtained from the ranked entries of Δ**B** from pooled results for the 50 network replicates. Two lists of ranked network edges were obtained from GENIE3: one for **B** and the other for 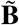. For each cutoff value of the rank, we obtain an adjacency matrix **A** from **B** as follows: the (*i, j*)th entry of **A** *a_ij_* = 1 if *b_ij_* is above the cutoff value, and otherwise *a_ij_* = 0. Similarly, another adjacency matrix **Ã** was obtained from 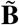. Then, the PR curve for changed edges detected by GENIE3 was obtained from **A** – **Ã**, again from pooled results for the 50 network replicates.

Figures 3 and 4 depict the PR curves of ProGAdNet, Lasso, and GENIE3 for measurement noise variance *σ*^2^ = 0.05 and 0.5, respectively. The number of samples varies from 50, 100, 200 to 300. It is seen from Figure 3 that our ProGAdNet offers much better performance than Lasso and GENIE3. When the noise variance increases from 0.05 to 0.5, the performance of all three algorithms degrades, but our ProGAdNet still outperforms Lasso and GENIE3 considerably, as shown in Figure 4. Table 1 lists AUPRs of ProGAdNet, Lasso and GENIE3, which again shows that our ProGAdNet outperforms Lasso and GENIE3 consistently at all sample sizes.

**Figure 3.**
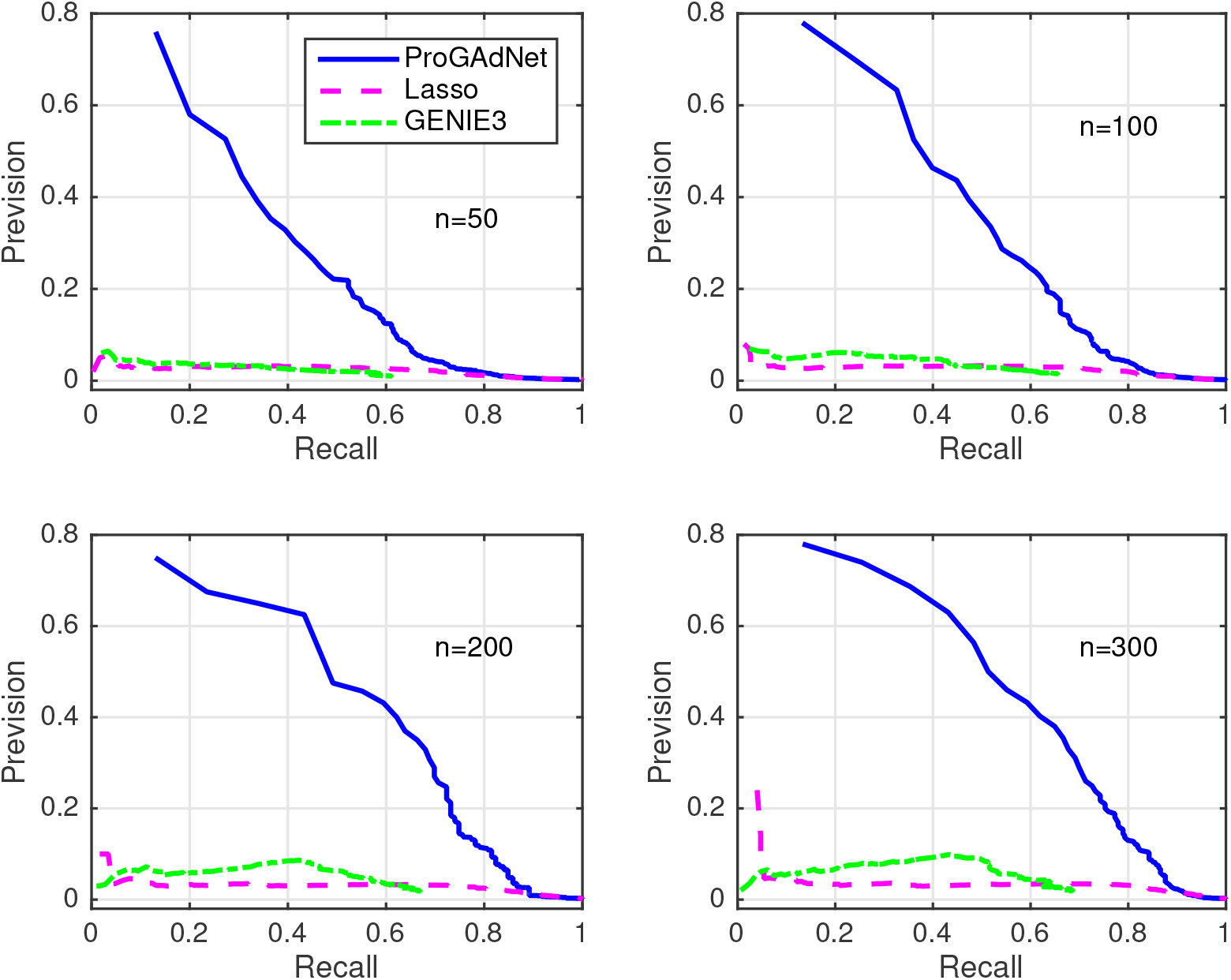
Precision-recall curves for ProGAdNet, Lasso, and GENIE3 in detecting changed edges of simulated gene networks. Variance of the measurement noise is σ^2^ = 0.05, and sample size *n*=50, 100, 200, and 300.

**Figure 4.**
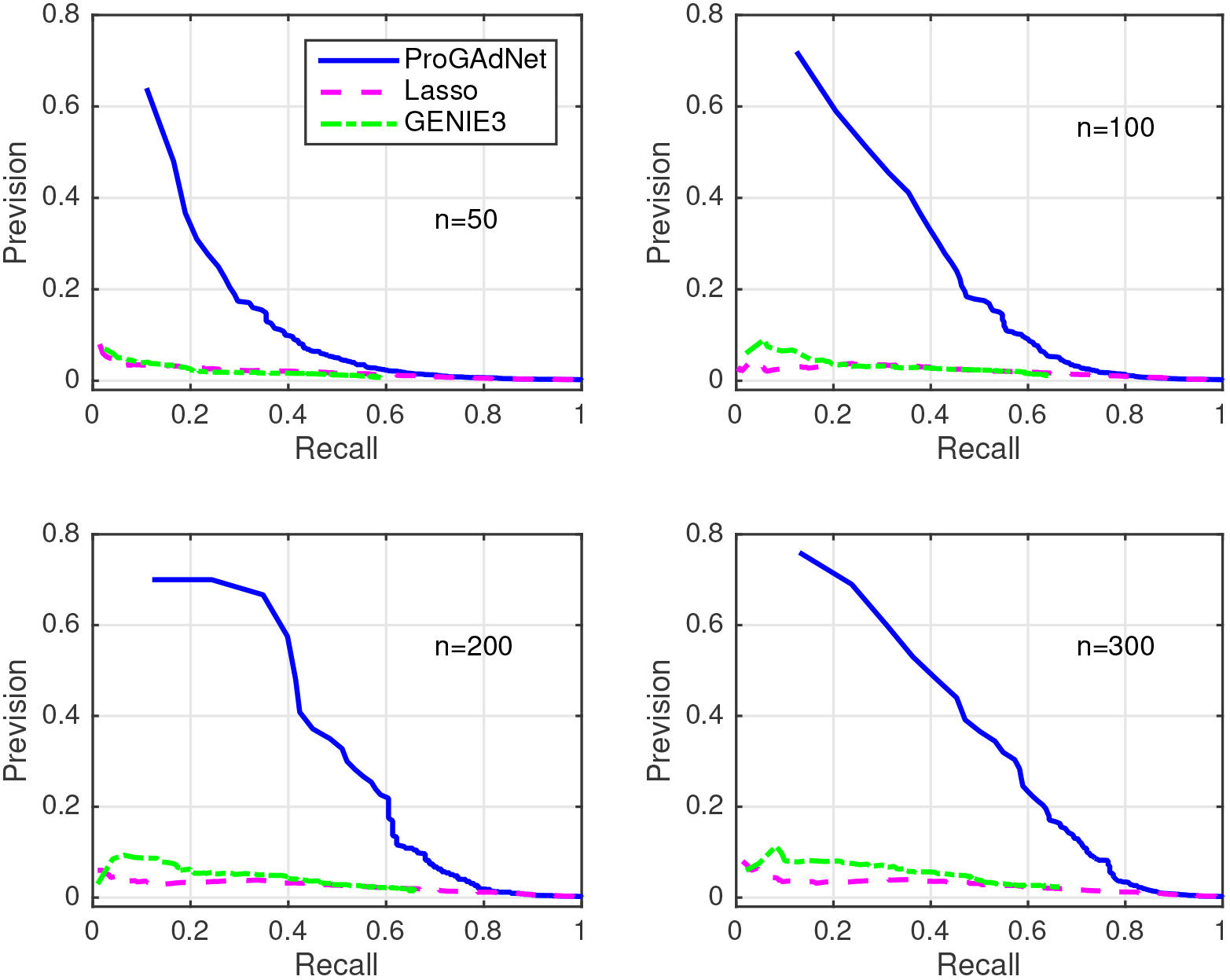
Precision-recall curves for ProGAdNet, Lasso, and GENIE3 in detecting changed edges of simulated gene networks. Variance of the measurement noise is σ^2^ = 0.5, and sample size *n*=50, 100, 200, and 300.

**Table 1.** AUPRs of ProGAdNet, Lasso, and GENIE3 for detecting the changed edges of simulated gene networks

| # samples | $\sigma^2 = 0.05$ | | | $\sigma^2 = 0.5$ | | |
| --- | --- | --- | --- | --- | --- | --- |
|  | ProGAdNet | Lasso | GENIE3 | ProGAdNet | Lasso | GENIE3 |
| 50 | 0.206 | 0.023 | 0.018 | 0.106 | 0.018 | 0.014 |
| 100 | 0.288 | 0.025 | 0.028 | 0.202 | 0.021 | 0.022 |
| 200 | 0.356 | 0.030 | 0.039 | 0.280 | 0.024 | 0.031 |
| 300 | 0.380 | 0.031 | 0.044 | 0.289 | 0.026 | 0.038 |

### Real Data Analysis

We next use the ProGAdNeT algorithm to analyze RNA-seq data of breast tumors and normal tissues. In The Cancer Genome Atlas (TCGA) database, there are RNA-seq data for 1,101 breast invasive carcinoma (BRCA) samples and 113 normal tissues. Although the TCGA database contains RNA-seq data of a number of other cancers, only the BRCA dataset has more than 100 normal tissue samples, and all other datasets contain less than 60 normal tissue samples. Since a small number of samples may compromise the accuracy of network inference, we only analyzed the BRCA dataset. The RNA-seq level 3 data for 113 normal tissues and their matched BRCA tumors were downloaded. The TCGA IDs of these 226 samples are given in Additional file 1. The scaled estimates of gene expression levels in the dataset were extracted, and they were multiplied by 10^6^, which yielded transcripts per million (TPM) value of each gene. The batch effect was corrected with the removeBatchEffect function in the Limma package [44] based on the batch information in the TCGA barcode of each sample (the “plate” field in the barcode). The RNA-seq data include expression levels of 20,531 genes. Two filters were used to obtain informative genes for further network analysis. First, genes with their expression levels in the lower 30 percentile were removed. Second, the coefficient of variation (CoV) was calculated for each of the remaining genes, and then genes with their CoVs in the lower 70 percentile were discarded. This resulted in 4,310 genes, and their expression levels in 113 normal tissues and 113 matched tumor tissues were used by the ProGAdNet algorithm to jointly infer the gene networks in normal tissues and tumors, and then to identify the difference in the two gene networks.”

Since small changes in *b_ji_* in the network model (1) may not have much biological effect, we regarded the regulatory effect from gene *j* to gene *i* to be changed using the following two criteria rather than the simple criterion 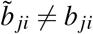. The first criterion is 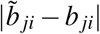 ≥ min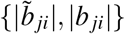, which ensures that there is at least one-fold change relative to min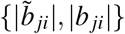. However, when one of 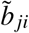 and *b_ji_* is zero or near zero, this criterion does not filter out very small 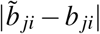. To avoid this problem, we further considered the second criterion. Specifically, nonzero 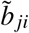 and *b_ji_* for all *j* and *i* were obtained, and the 20 percentile value of all 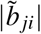 and *|b_ji_|, T*, was found. Then, the second criterion is max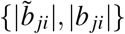 ≥ *T*. As in computer simulations, the stability selection was employed to identify network changes reliably. As the number of genes, 4,310, is quite big, it is time consuming to repeat 100 runs per *λ*_1_ and *λ*_2_ pair. To reduce the computational burden, we used five-fold cross validation to choose the optimal values of *λ*_1_ and *λ*_2_ based on the two-SE rule used in computer simulation, and then performed stability selection with 100 runs for the pair of optimal values. Note that stability selection at an appropriate point of hyperparameters is equally valid compared with that done along a path of hyperparameters [39]. The threshold for Δ*r_ij_* for determining network changes as described in the Method section was chosen to be *c* = 0.9.

Our network analysis with ProGAdNeT identified 268 genes that are involved in at least one changed edge. Names of these genes are listed in supplementary file 2. We named the set of these 268 genes as the dNet set. To assess whether the dNet genes relate to the disease status, we performed gene set enrichment analysis (GSEA) with the C2 gene sets in the molecular signatures database (MSigDB) [45, 46]. C2 gene sets consist of 4,738 gene sets that include pathways in major pathway dabases such as KEGG [47], REACTOME [48], and BIOCARTA [49]. After excluding gene sets with more than 268 genes or less than 15 genes, 2,469 gene sets remained. Searching over the names of these 2,469 gene sets with key words “breast cancer”, “breast tumor”, “breast carcinoma” and “BRCA” identified 139 gene sets that are related to breast cancer. Using Fisher’s exact test, we found that 78 of 2,469 C2 gene sets were enriched in the dNet gene set at a FDR of < 10^−4^. The list of the 78 gene sets is in Additional file 3. Of these 78 gene sets, 24 are among the 139 breast cancer gene sets, which is highly significant (at Fisher’s exact test p-value 2.3 × 10^−9^). The top 20 enriched gene sets are listed in Table 2. As seen from names of these gene sets, 11 of the 20 gene sets are breast cancer gene sets, and 7 sets are related to other types of cancer. These GSEA results clearly show that the dNet gene set that our ProGAdNet algorithm identified is very relevant to the cancer disease status.

**Table 2.** Top 20 MSigDB C2 gene sets that are enriched in the dNet gene set.

| Gene sets | q-value |
| --- | --- |
| SMID_BREAST_CANCER_LUMINAL_A_UP | 1.55E-27 |
| NAKAYAMA_SOFT_TISSUE_TUMORS_PCA2_DN | 1.73E-20 |
| TURASHVILI_BREAST_DUCTAL_CARCINOMA_VS_DUCTAL_NORMAL_DN | 1.40E-16 |
| SMID_BREAST_CANCER_RELAPSE_IN_LUNG_DN | 1.40E-16 |
| POOLA_INVASIVE_BREAST_CANCER_UP | 1.25E-12 |
| SMID_BREAST_CANCER_RELAPSE_IN_BONE_UP | 2.35E-12 |
| TURASHVILI_BREAST_DUCTAL_CARCINOMA_VS_LOBULAR_NORMAL_DN | 2.72E-12 |
| VECCHI_GASTRIC_CANCER_ADVANCED_VS_EARLY_UP | 3.04E-12 |
| MCLACHLAN_DENTAL_CARIES_UP | 3.04E-12 |
| POOLA_INVASIVE_BREAST_CANCER_DN | 3.04E-12 |
| TURASHVILI_BREAST_LOBULAR_CARCINOMA_VS_DUCTAL_NORMAL_DN | 9.32E-12 |
| CROMER_TUMORIGENESIS_UP | 1.26E-11 |
| TURASHVILI_BREAST_LOBULAR_CARCINOMA_VS_LOBULAR_NORMAL_UP | 1.52E-11 |
| DOANE_BREAST_CANCER_ESR1_UP | 1.52E-11 |
| LIEN_BREAST_CARCINOMA_METAPLASTIC_VS_DUCTAL_DN | 2.28E-11 |
| SABATES_COLORECTAL_ADENOMA_DN | 3.70E-09 |
| JAEGER_METASTASIS_DN | 4.31E-09 |
| WALLACE_PROSTATE_CANCER_RACE_UP | 4.46E-09 |
| ANASTASSIOU_MULTICANCER_INVASIVENESS_SIGNATURE | 4.68E-09 |
| KORKOLA_TERATOMA | 7.24E-09 |

## Conclusion

In this paper, we developed a very efficient algorithm, named ProGAdNet, for inference of two gene networks based on gene expression data under two different conditions, which were further used to identify differential changes in the network. Computer simulations showed that our ProGAdNet offered much better inference accuracy than existing algorithms. Analysis of a set of RNA-seq data of breast tumors and normal tissues with ProGAdNet identified a set of genes involved in differential changes of the gene network. A number of gene sets of breast cancer or other types of cancer are significantly enriched in the identified gene set, which shows that the identified gene set is very informative about the disease status of the tissues. As gene network rewiring occurs frequently under different molecular context, our ProGAdNet algorithm provides a valuable tool for identifying changed gene-gene interactions.

## Availability of data and materials

The software package implementing the ProGAdNet algorithm and computer simulations is available upon request. The RNA-Seq data of 113 breast tumors and 113 normal tissues used in differential gene network analysis can be downloaded from the website of Gnomic Data Commons at https://Gd.cancer.gov/ using the TCGA sample IDs in Additional file 1.

## Author’s contributions

XC conceived the idea and supervised the project. XC, CW, and FG designed the algorithm. CW performed computer simulation with linear regression models and analysis of real data. FG performed computer simulation with gene networks. GBG contributed to the development of the algorithm and computer simulations. GD contributed to real data analysis. XC, CW, and FG wrote the manuscript. All authors read and approved the final manuscript.

## Additional files

Additional file 1: the list of the TCGA IDs of 113 breast tumors and 113 normal tissue samples.

Additional file 2: the list of 268 genes that are involved in at least one changed network edges identified from the gene expression data of breast tumor and normal tissues.

Additional file 3: the list of 78 MSigDB C2 gene sets that are significantly enriched in the gene sets in Additional file 2.

